# Microglia drive diurnal variation in susceptibility to inflammatory blood-brain barrier breakdown

**DOI:** 10.1101/2024.04.10.588924

**Authors:** Jennifer H. Lawrence, Asha Patel, Melvin W. King, Collin J. Nadarajah, Richard Daneman, Erik S. Musiek

## Abstract

The blood-brain barrier (BBB) is critical for maintaining brain homeostasis but is susceptible to inflammatory dysfunction. Permeability of the BBB to lipophilic molecules shows circadian variation due to rhythmic transporter expression, while basal permeability to polar molecules is non-rhythmic. Whether daily timing influences BBB permeability in response to inflammation is unknown. Here, we induced systemic inflammation through repeated lipopolysaccharide (LPS) injections either in the morning (ZT1) or evening (ZT13) under standard lighting conditions, then examined BBB permeability to a polar molecule, sodium fluorescein. We observed clear diurnal variation in inflammatory BBB permeability, with a striking increase in paracellular leak across the BBB specifically following evening LPS injection. Evening LPS led to persisting glia activation and inflammation in the brain that was not observed in the periphery. The exaggerated evening neuroinflammation and BBB disruption were suppressed by microglial depletion or through keeping mice in constant darkness. Our data show that diurnal rhythms in microglial inflammatory responses to LPS drive daily variability in BBB breakdown and reveals time-of-day as a key regulator of inflammatory BBB disruption.

## Introduction

Activation of the innate immune response in the brain drives neuroinflammation which is a critical component of many neurological disorders. Innate immune activation strongly influences many aspects of disease pathogenesis in age-related neurodegenerative disorders, including Alzheimer’s Disease, as well as in more acute insults such as infection or stroke (1–4). In mice, inflammatory responses to a variety of inflammogens, including the toll-like receptor 4 agonist lipopolysaccharide (LPS), show striking time-of-day effects (5–8). Mice treated with systemic LPS in the evening have increased peripheral inflammation and mortality than those treated in the morning (5,9,10) . The circadian clock system plays a role in regulating innate immune responses (11) and this time-of-day difference in LPS-mediated mortality can be ablated by disrupting the core circadian clock via deletion of the key clock gene *Bmal1* (5). However, LPS-induced inflammatory responses in the periphery, as well as animal mortality, are governed not only by the circadian clock, but also a host of other factors including exposure to light and time of feeding, and do not seem to rely on the circadian function of monocytes (5,6,11,12). Moreover, while diurnal variation in the severity of innate immune activation has been demonstrated in peripheral organs, it is unknown if similar time-of-day variations in the severity of inflammation occur in the brain, and if so, which cells might regulate this phenomenon.

One consequence of neuroinflammation is dysfunction of the blood-brain barrier (BBB). The BBB is critical for restricting certain blood-borne molecules from the brain, helping protect the central nervous system (CNS) from circulating neurotoxic and neuroinflammatory molecules as well as peripheral immune cells (13–15). The BBB is a multicellular structure, consisting of endothelial cells, mural cells (pericytes and smooth muscle cells), and astrocytes - collectively referred to as the neurovascular unit (NVU). Outside of the NVU, microglia and neurons also impact BBB function (16,17). This selectively permeable barrier is formed by a series of distinct cellular properties within the NVU, including continuous non-fenestrated vessels, decreased pinocytosis, the presence of junctional proteins, and additional regulatory elements that allow for the selective movement of solutes between the blood and CNS (18). In the setting of peripheral or central inflammation, the BBB can become more permeable due to transporter dysfunction, breakdown of endothelial cell tight and adherens junctions, and increases in transcytosis (19,20). While BBB permeability to lipophilic molecules has been reported and is dependent on circadian expression of specific transporter proteins such as p-glycoprotein (17,21), the BBB does not show daily rhythms in permeability to polar molecules under basal conditions (17,22–24). However, it is unknown how disease processes such as inflammation might influence time-of-day variations in BBB permeability, as many cellular functions can gain diurnal rhythms in the setting of pathology (25–28).

Here, we have investigated how the neuroinflammatory response and BBB permeability to polar molecules (termed inflammatory BBB breakdown) are influenced by innate immune activation (exposure to intraperitoneal LPS injection) at different times of day. We observed that both inflammatory gene expression in the brain and BBB breakdown are more severe following LPS exposure in the evening (ZT13) as compared to morning (ZT1). This diurnal variation appears to depend on light exposure and is characterized by persisting astrocyte and microglial activation and cytokine production after evening LPS exposure which is not observed in the morning and is not present in peripheral organs. We find that exaggerated neuroinflammatory BBB disruption in the evening occurs independently of rhythms in peripheral inflammation and is instead intrinsic to CNS-resident immune cells, particularly microglia.

## Results

### Evening LPS exposure causes persisting neuroinflammation and gliosis

We first sought to determine if exposure to an inflammatory stimulus at different times of day can differentially impact CNS inflammation. To induce robust neuroinflammation, we gave daily intraperitoneal (i.p.) injections of 2 mg/kg LPS or PBS control to 2-month-old male C57bl/6 mice for two days in a row at either 7am or 7pm. In a 12-hour light cycle, these times correspond to 1 hour after the onset of light (ZT1) and 1 hour after the lights turn off (ZT13). We chose these times because they roughly correspond to the nadir and peak of diurnal behavioral rhythms, as well as circadian gene expression in most organs (29). In models of systemic LPS administration, peripheral cytokine mRNA levels peak between 3-6 hours post injection (hpi) (30–33), so we collected peripheral and CNS tissue 6- and 24-hours after the final ZT1 or ZT13 dose of LPS (6 hour time points were 1pm (ZT7) or 1am (ZT17)), and examined inflammatory mRNA expression of a subset of transcripts by qPCR. We chose lung tissue to serve as a marker of peripheral inflammation due to its known circadian rhythmicity in LPS response with minimal impact of time of feeding (6,11). We saw no diurnal differences in lung cytokine production or percent weight loss between the ZT1 and ZT13-injected groups at either 6 or 24 hours post-LPS (Figure S1A, B). Additionally, we saw no diurnal differences in inflammatory gene expression in the liver at 6 hpi or the spleen at 24 hpi (Figure S1C, D); altogether, these data indicate that, in this particular model of systemic inflammation, there are no major diurnal variations in peripheral inflammatory response.

In cortical tissue, LPS strongly induced inflammatory gene expression at 6 hpi, but diurnal differences were not evident. However, at 24 hpi, several inflammatory genes (*Tnfa*, *Cxcl5*, and *Nos2*) showed increased expression in the Z13 LPS group (Figure 1A). When comparing the 6 hpi to 24 hpi gene expression datasets, there is a sharp reduction in cortical inflammation at 24 hpi in all groups though this resolution of inflammatory gene expression is dampened in the ZT13 LPS group (Figure 1A, B). Conversely, markers of gliosis remain persistently elevated at both 6 and 24 hpi, although the microglial activation marker *Cybb* gained a diurnal difference 24 hours after LPS (Figure 1A, B). We next quantified micro- and astrogliosis by immunohistochemistry in the same groups of mice (ZT1 or ZT3 LPS) sacrificed at 6 or 24 hpi. At 6 hpi there were no diurnal effects on microgliosis as measured by IBA1 percent area in the hippocampus, although there was a treatment effect at only ZT13 (Figure 1C). Similarly, using GFAP as a marker for astrogliosis in the hippocampus, there were no LPS or time effects found at 6 hpi (Figure 1C). However, there was a significant diurnal effect on IBA1 and GFAP immunoreactivity 24 hours post LPS, with both microglia and astrocytes showing increased activation in the ZT13 LPS group (Figure 1D). These data suggest that there may be decreased resolution of inflammation and microgliosis in mice treated with LPS at ZT13 compared to those at ZT1.

**Figure 1:**
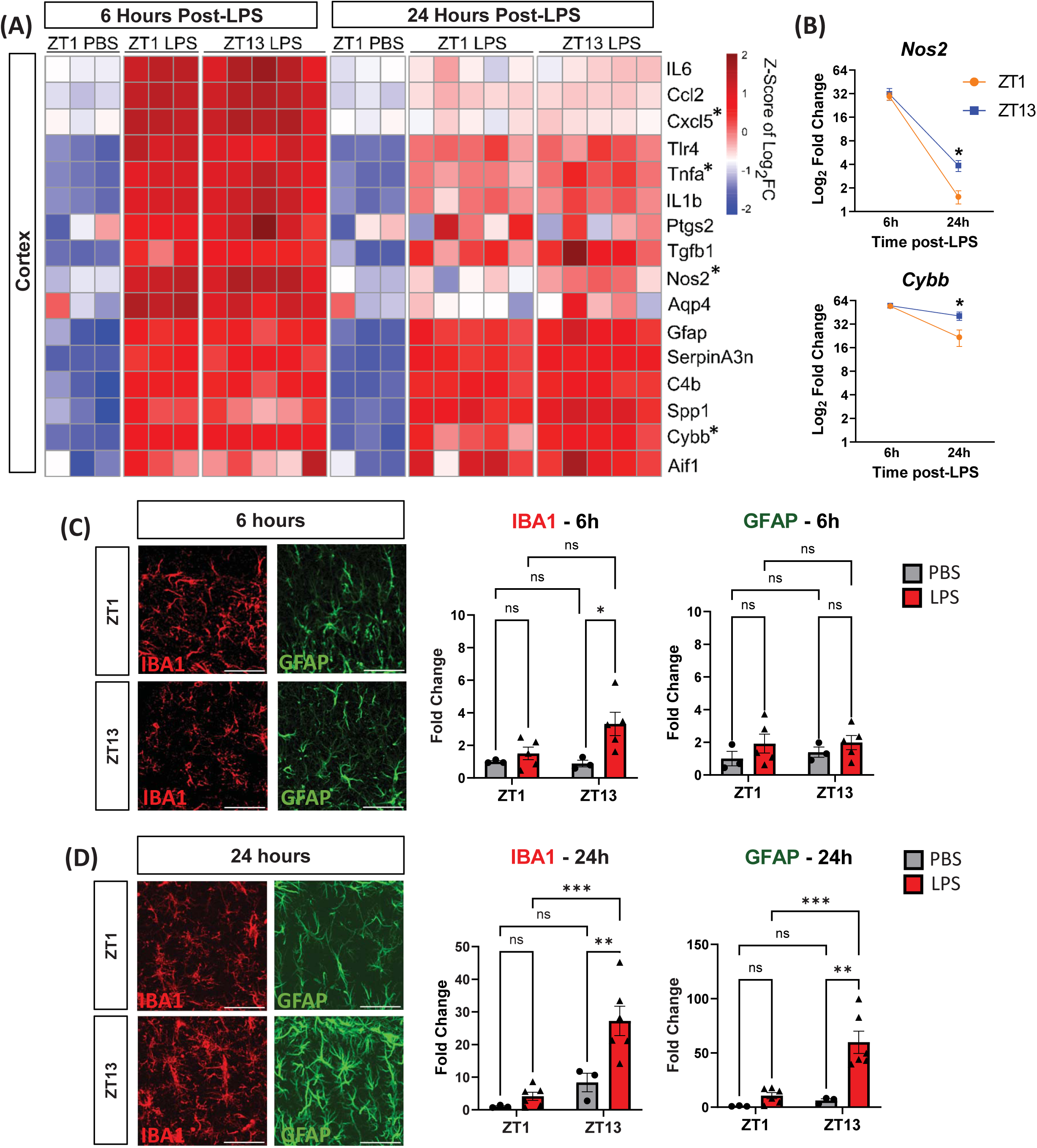
Evening LPS causes persisting inflammation and gliosis. (A) Heatmap of selected gene expression in mouse cortex 6- vs 24-hours after final LPS injection at ZT1 or ZT12. Gene expression is normalized to ZT1 PBS 6 hpi group, Transcripts with significant TOD effect at 24 hpi are denoted with *. Scale is Z-Score of Log_2_FC. (B) *Nos2* and *Cybb* graphed across time (significant TOD effects are denoted with *). (C) Representative images of IBA1 and GFAP after 6h of PBS or LPS treatment with quantification of percent area in the hippocampus 6 or 24 hours post-LPS, normalized to ZT1 PBS group. (D) Representative images of IBA1 and GFAP after 24h of PBS or LPS treatment with quantification of percent area in the hippocampus 6 or 24 hours post-LPS, normalized to ZT1 PBS group. n = 3-5 mice per group. For 6 hr timepoint, only main effect of LPS was significant for IBA1, while in 24 hrs timepoint main effect of LPS and interaction are significant for both proteins, and post-hoc test results are shown. *p<0.05, **p<0.01, ***p<0.005, ****p<0.001. Scale bars in (C) and (D) are 50um.

Rhythms in immune response may confer an evolutionary advantage by ensuring maximum immunity during times of day when animals are at the greatest risk of injury or infection (34). Since i.p. LPS induces robust systemic inflammation, it is important to disentangle the role of rhythms in peripheral and central inflammation. We found no diurnal variation in body weight loss, mortality, or peripheral cytokine production across multiple organs and time points, indicating there is no diurnal variation in peripheral LPS response - at least in this two-hit LPS model. While brain endothelial cells (BECs) are TLR4^+^ and can directly bind to LPS, LPS protein is not found in the brain parenchyma after systemic administration (30). This indicates that rather than interacting directly with CNS-resident cells, LPS most likely acts indirectly on peripheral TLR4^+^ cells – including leukocytes as well as BECs – producing cytokines which can readily cross the BBB. Additionally, existing circadian sequencing databases have found that there are no circadian rhythms in *TLR4* expression in BECs – suggesting that rhythmic neuroinflammatory responses to LPS occurs through CNS-resident cells (21). Specifically, increased astro- and microgliosis in the evening indicate there is diurnal regulation of glial immune responses to these peripherally derived cytokines, persisting up to 24 hours post-LPS.

### Persisting neuroinflammation after evening LPS specifically enhances BBB-related gene expression

To fully characterize neuroinflammation 24h post-LPS, we conducted bulk RNAseq on cortex samples from mice treated with i.p. LPS (or PBS control) at ZT1 and ZT13. While previous studies have characterized the brain inflammatory profile induced by LPS, none have considered diurnal variations in immune response to LPS exposure (35–37). When comparing only LPS-treated mice between ZT1 and ZT13, we identified 209 differentially expressed genes (DEGs; 181 upregulated and 28 downregulated, adjusted P<0.05) in the ZT13 LPS compared to ZT1 LPS group (Figure 2A). Many of these genes upregulated in the evening were pro-inflammatory mediators (*Nos2, Mmp8, Stab1, Cd14*). Since the majority of DEGs were upregulated at ZT13, we next asked if LPS treatment induced more transcripts at ZT13 than at ZT1 by examining PBS vs. LPS at ZT1 and separately analyzing PBS vs. LPS at ZT13, then comparing the DEG lists. There were 3287 DEGs (1761 upregulated and 1526 downregulated) in LPS compared to PBS treated mice at ZT1 and 5398 DEGs (2756 upregulated and 2642 downregulated) in LPS compared to PBS treated mice at ZT13. Comparisons between these two datasets found that there were 2661 DEGs uniquely found in mice treated at ZT13, with only 550 DEGs unique to ZT1, and an additional 2737 DEGs found at both times (Figure 2B).

**Figure 2:**
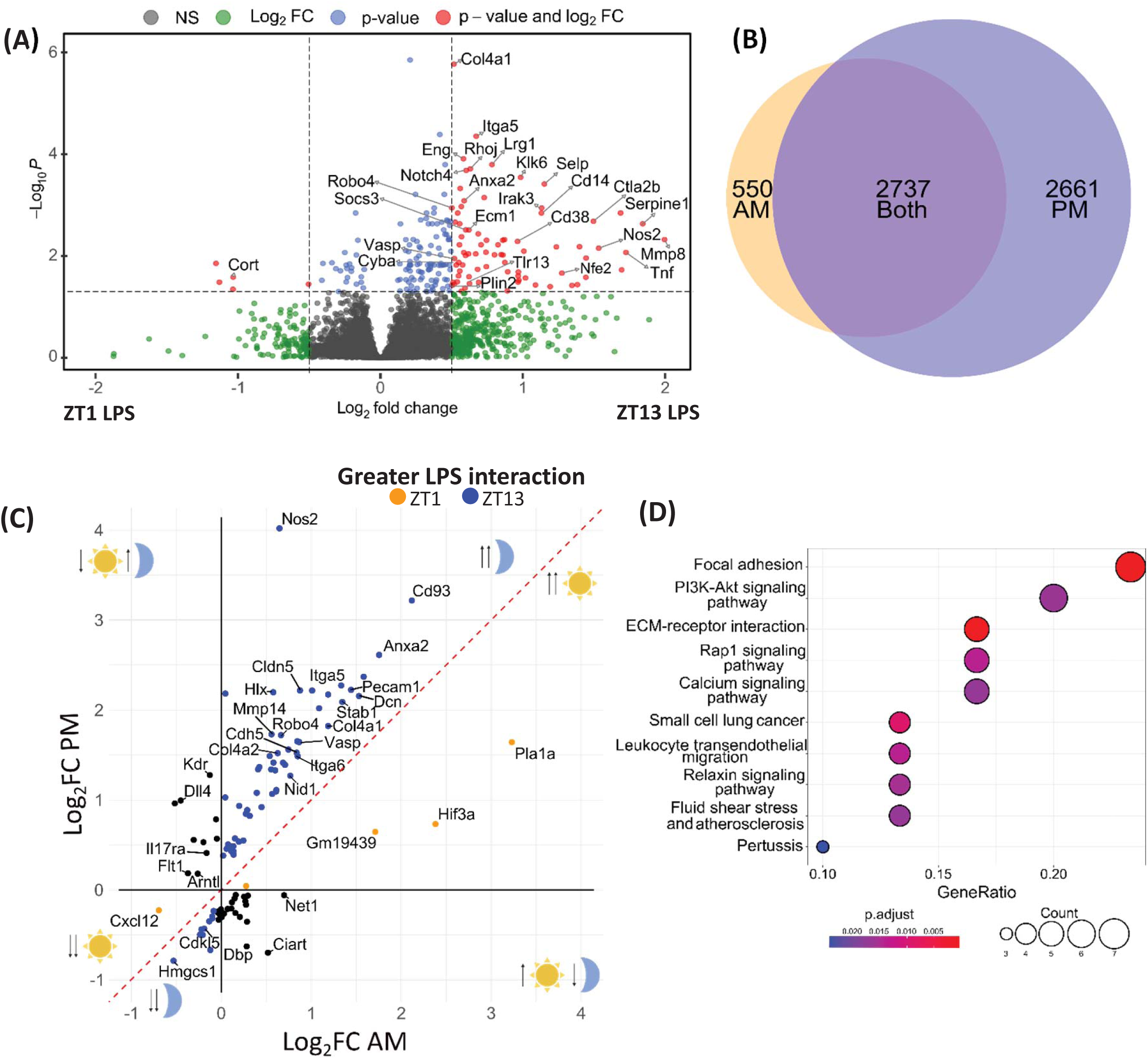
Persisting neuroinflammation after evening LPS exposure specifically enhances BBB-related gene expression. (A) Volcano plot of differentially expressed genes (DEGs) from bulk RNAseq in LPS-treated mouse cortex, comparing ZT1 vs. ZT13 treatment time. Red dots indicate transcripts with adjusted P value <0.05 and fold change >1.75 fold, and right upper area indicates higher expression in mice treated at ZT13 (B) Venn diagram showing overlap of DEGs upregulated after LPS in the ZT1 vs. ZT13 datasets. (C) XY plot of genes with a significant TOD and treated interaction by 2-way ANOVA. Log fold change (PBS vs. LPS) in the AM (ZT1) group is shown on X axis, and in the PM (ZT13) group on Y axis. Cartoons indicate the response of genes in that area to LPS (up or down arrow) in the AM (sun) or PM (moon). Genes shown above the red dotted line and right of the Y-axis were all upregulated in the PM, with *Nos2* being the highest fold change. (D) ORA dotplot of the top dysregulated KEGG pathways using the full gene list from D. n = 3-5 mice per group

We next used a 2-way ANOVA to examine statistical interaction between our two variables, time-of-day (TOD, ZT1 vs. ZT13) versus treatment (PBS vs. LPS). This method allowed us to control for baseline rhythms in gene expression in the PBS control groups. We then plotted only the DEGs with a significant TOD x LPS interaction on axes representing their log_2_fold change in AM (x-axis) versus PM (y-axis). This showed us that DEGs which are upregulated in response to LPS tend to be further upregulated when treated with LPS in the PM (Figure 2C). KEGG pathway analysis of all DEGs with a significant TOD x treatment interaction found dysregulation of several vascular related functions - focal adhesion and fluid shear stress and atherosclerosis - as well as inflammatory pathways, including leukocyte transendothelial migration and ECM-receptor interaction (Figure 2D). Interestingly, many of the genes with a TOD x treatment interaction are involved in BBB function, including upregulation of the TJ- and AJ-associated proteins *Cldn5* and *Cdh5* (Figure 2C). Additionally, there was a diurnal increase in expression of the vasodilator-stimulated protein (*Vasp*) and inducible nitric oxide synthase (*Nos2*) genes, both of which have been implicated in BBB dynamics (Figure 2C) (38,39). Altogether, while we saw significant changes in LPS-induced gene expression at both the ZT1 and ZT13 time points, BBB-associated genes were among the most significantly dysregulated pathways when considering the interaction of time and treatment.

### Evening LPS exposure disrupts endothelial cell morphology

Since BBB-associated gene pathways were highly dysregulation in response to both time-of-day and LPS, we used transmission electron microscopy (TEM) to image cortical capillary ultrastructure (Figure S2A). Solutes can cross the BBB in a myriad of ways including passive paracellular or transcellular diffusion as well as adsorptive-, carrier-, or receptor-mediated transport. Inflammatory BBB breakdown occurs primarily through either disruption of endothelial tight junctions (TJs) or an increase in transcytosis (40). While we saw no gross disruption of the neurovascular unit (NVU), there was significant diurnal variation in vasodilation, with an increase in capillary diameter in mice treated with LPS at ZT13 as compared to ZT1 (Figure 3A, B) – consistent with the increase in *Nos2* and *Vasp* seen in our RNAseq dataset. Despite known circadian rhythms in *Claudin5* gene expression (41), we measured the average tight junction (TJ) length per vessel and saw no change based on time or treatment (Figure 3A, C). Although, we did note select disrupted TJs imaged in the LPS treatment group (Figure S2B).

**Figure 3:**
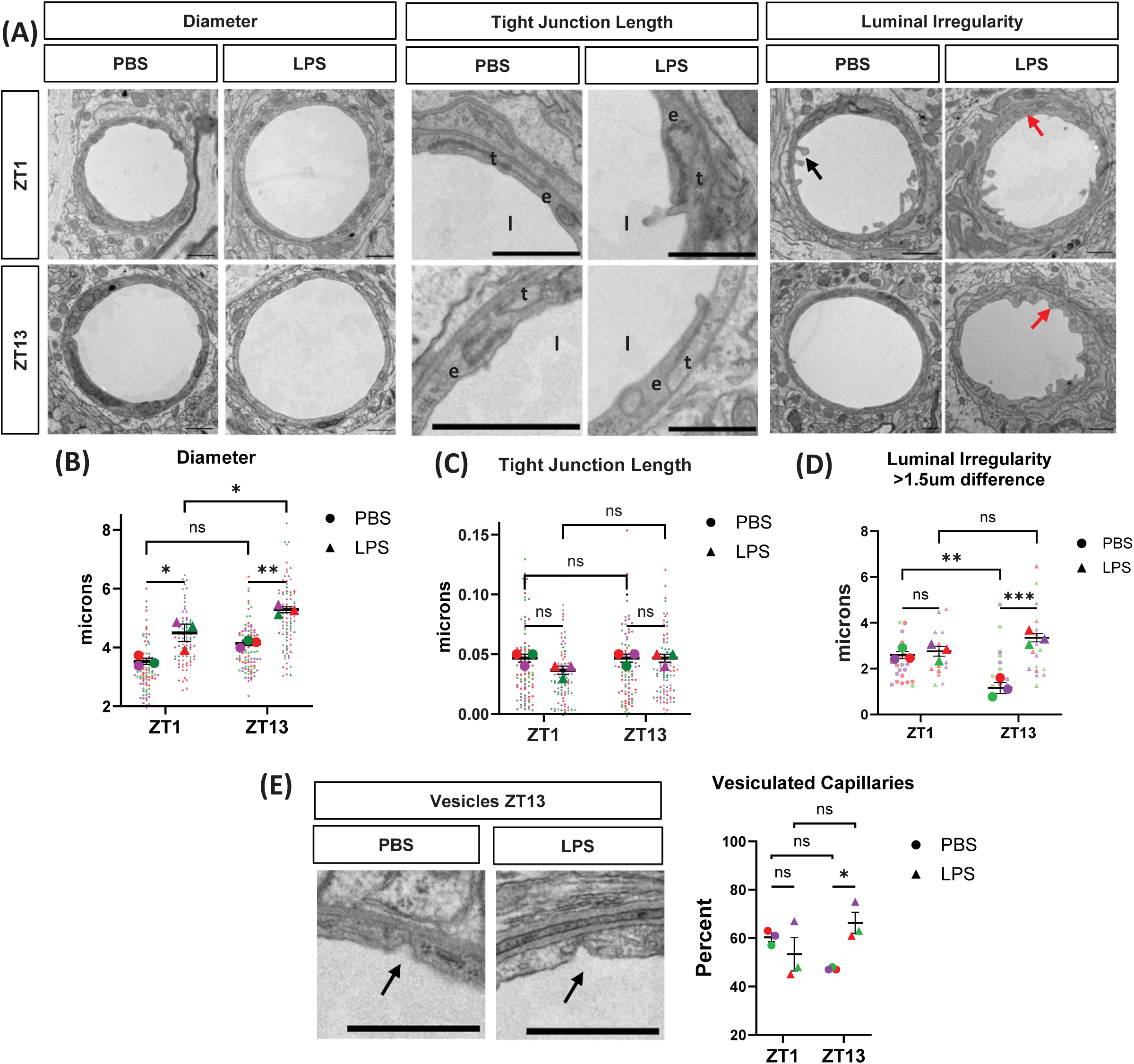
Evening LPS exposure disrupts endothelial cell morphology. (A) Representative images of diameter, tight junction length (e = endothelial cell, l = lumen, t = tight junction), and luminal irregularity (arrows: black showing blebbing at luminal surface, red showing plasma/basement membrane disruption). (B) Quantification of diameter length across cortical capillaries; main effect of LPS was significant, but interaction was not by 2-way ANOVA. (C) Quantification of average tight junction length per capillary; no effects were significant by 2-way ANOVA. (D) Quantification of luminal irregularity; main effect of TOD and Interaction were significant by 2-way ANOVA, and post-hoc test is shown. (E) Inclusion criteria for vesicles (arrow indicating invaginating vesicles) and quantification of percentage of capillaries that contain one or more vesicles. Interaction was significant by 2-way ANOVA, and post-hoc test is shown. For all graphs, N = 3 mice per group, 28-32 capillaries per mouse. Average for each mouse is shown in large dots, while technical replicates shown as smaller dots in similar color per mouse; Scale bar = 1 micron.

Previous TEM studies have also found that LPS induces luminal plasma membrane ruffling, a mechanism of non-receptor mediated pinocytosis used by bacteria and viruses to increase infection of epithelial cells (20,42,43). We found a significant increase in severe luminal irregularity in LPS treated mice at ZT13 only (Figure 3A, D). To confirm that the increase in plasma membrane disruption seen in ZT13 LPS treated mice is indicative of increased transcytosis, we next quantified the number of vesicles per capillary, only counting actively invaginating vesicles from the luminal side. We found a significant increase in the percentage of vessels containing at least one vesicle in LPS treated mice at ZT13 only (Figure 3E). Altogether, these ultrastructural abnormalities indicate evening LPS exposure induces increased vasodilation as well as a disrupted endothelial cell morphology that is consistent with increased paracellular or transyctotic particle uptake.

### Time-of-day (TOD) strongly regulates inflammatory BBB breakdown

To determine if the ZT13 LPS-induced endothelial cell morphology disruption and gene dysregulation lead to BBB breakdown, we administered systemic tracers and quantified the amount found in the hippocampus, cortex, and cerebellum - all normalized to the amount found in the serum (Figure 4A). First, in order to identify gross size exclusion properties in LPS-induced BBB breakdown, we examined BBB permeability to two polar tracers of varying size – with mice treated only at ZT13, the time at which there is greatest LPS-induced inflammation and mortality (Figure 2C) (5,44). Evans Blue (EB) is a relatively small azo dye (961 Da) that binds with high affinity to serum albumin, creating a high molecular weight (mw) tracer of 69 kDa (45). Conversely, sodium fluorescein (NaFL) is a small mw (376 Da) inert fluorescent molecule that weakly binds to endogenous proteins and has no known endothelial cell transporters, allowing it to serve as a sensitive BBB permeability tracer (46). Neither phosphate buffer saline (PBS) control nor LPS-treated mice showed any visible leak of EB, indicating there is no gross hemorrhaging or permeability to larger molecules in this model of inflammation (Figure 4B). While there was no NaFL seen in the brain at baseline in PBS control mice, there was robust NaFL leak in ZT13 LPS treated mice (Figure 4B). Thus, all further tracer experiments were conducted using NaFL.

**Figure 4:**
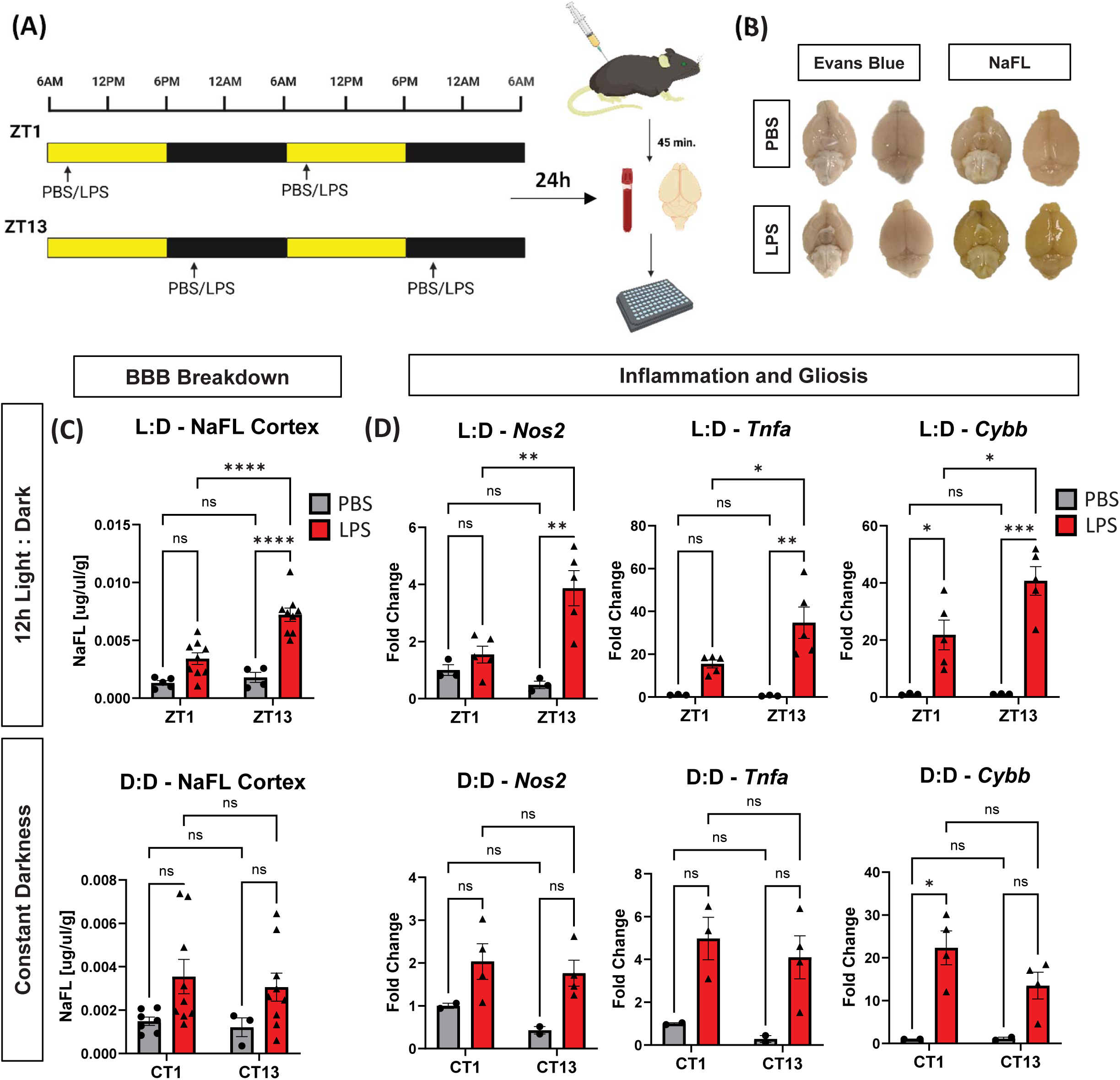
Light drives diurnal variation in inflammatory BBB breakdown. (A) Schematic of the 2-hit LPS experimental paradigm used for diurnal BBB permeability experiments using sodium fluorescein (NaFL), with lights-on (ZT0) occurring at 6am and lights-off (ZT12) occurring at 6pm. (B) Representative images of dorsal and ventral view of whole brains from mice injected with PBS and LPS as in A, then given either Evans Blue or NaFL tracer. (C) Quantification of diurnal differences in LPS-induced NaFL leak in the cortex for mice kept in L:D or D:D; CT1 is 7am and CT13 is 7pm. n = 3-10 mice per group. For L:D data, both main effects of treatment and time of day, as well as interaction, were significant by 2-way ANOVA. Post-hoc test P values are shown. (D) qPCR data from cortex of L:D and D:L PBS or LPS treated mice. n = 3-5 mice per group. Main effect of treatment and interaction were significant by 2-way ANOVA; post-hoc P values are shown. *p<0.05, **p<0.01, ***p<0.005, ****p<0.001.

We next examined the effect of time-of-day (TOD) of LPS exposure on BBB permeability. As expected, there was minimal permeability of NaFL in PBS-treated mice at either ZT1 or ZT13 (Figure 4C, S2C), supporting previous studies stating that basal permeability to polar molecules does not show diurnal variation (17,22). However, mice treated with LPS in the evening (ZT13) had significantly greater NaFL leak across multiple brain regions than those treated in the morning (ZT1), with the cerebellum having the greatest absolute leak and the cortex having the strongest TOD effect (Figure 4C, S2C). Of note, while LPS treatment at ZT1 only slightly increased BBB permeability to NaFL in the cortex and hippocampus, permeability was significantly increased in all brain regions after LPS exposure at ZT13.

Since constant darkness is known to abrogate the time-of-day effect of LPS-induced mortality (5), we next examined the effects of 24-hours of constant darkness (D:D) on LPS-induced BBB breakdown. We found that the diurnal differences in LPS-induced NaFL leak were ablated in all three brain regions in mice kept in D:D (Figure 4C, S2D). Overall, there was decreased NaFL leak at Circadian Time 13 (CT13), suggesting that light:dark conditions are required for diurnal increases in LPS-induced inflammatory BBB breakdown. We also found that D:D ablated time-of-day difference in gene expression associated with inflammation (*Nos2, Tnfa*) and microglial activation (*Cybb*), suggesting a general reduction of neuroinflammation (Figure 4D). Altogether, these data indicate that light:dark conditions are required for diurnal variation in neuroinflammation and inflammatory BBB breakdown.

While inflammatory BBB breakdown and circadian rhythms in barrier properties have both been explored independently, there is relatively little known about daily rhythms in neuroinflammation and BBB breakdown. Our current findings suggest that there are diurnal changes in the susceptibility of the BBB to inflammatory breakdown, beyond the known rhythms in transporter activity (17,22–24). Previous studies on LPS-induced mortality have found the phenomenon is light *dependent* and myeloid cell clock *independent*, as rhythms in mortality are ablated in mice kept in constant darkness (DD) and persist in mice deficient in the master circadian clock protein, BMAL1, in myeloid cells (5). Similarly, we found mice kept in constant darkness showed no diurnal variation in BBB breakdown or in markers of neuroinflammation or microgliosis. Of note, constitutive brain-specific deletion of BMAL1 has been reported to induce BBB breakdown at baseline, further complicating the study of cellular circadian rhythms in inflammatory BBB leakage (47).

### Microglia are required for LPS-induced neuroinflammatory BBB breakdown at ZT13

At baseline, capillary-associated microglia play an active role in maintaining BBB integrity. Ablating microglia (and other CSR1+ myeloid populations in the brain) with the CSFR1 antagonist Plexidartinib (PLX3397, referred to as PLX) alters vascular tone without changing pericyte coverage or astrocyte endfoot density (48,49). During inflammation, PLX treatment protects against LPS-induced inner blood retinal barrier leak in the eye but is deleterious during models of both ischemic stroke and micro-bubble induced hemorrhage in the CNS (16,50,51). Additionally, in our LPS model we found that the increase in gliosis seen at ZT13 was coupled with increased perivascular localization (within 8um of a blood vessel) of IBA1 but not GFAP (Figure S3A). To determine if microglia are required for LPS-induced BBB breakdown, we treated mice for 2 weeks with PLX prior to LPS administration. Since there was no significant increase in BBB permeability with LPS treatment at ZT1, we only treated PLX (or control chow) mice at the ZT13 time point. Of note, a previous study shows that PLX administration does not impact behavioral circadian rhythms in mice (52). As previously reported, we found that PLX treatment completely ablated IBA1^+^ cells (Figure 5B, C) (49) in both PBS and LPS treated groups without altering peripheral immune responses or sickness behaviors (Figure 5A) (53). Despite microglia being the primary TLR4^+^ CNS-resident cell, LPS-treated PLX mice had increased hippocampal GFAP (Figure 5B, C) as well as *Serapina3n* and *GFAP* upregulation (Figure S3B), suggesting that astrocyte activation in response to systemic LPS is independent of microglial function. However, despite these changes in astrocyte GFAP expression, PLX administration reduced LPS-induced *TNFa* and *Nos2* in cerebral cortex and completely protected against NaFL leak, indicating that microglia are required for neuroinflammation following peripheral LPS injection, as well as inflammatory BBB breakdown (Figure 5D, E).

**Figure 5:**
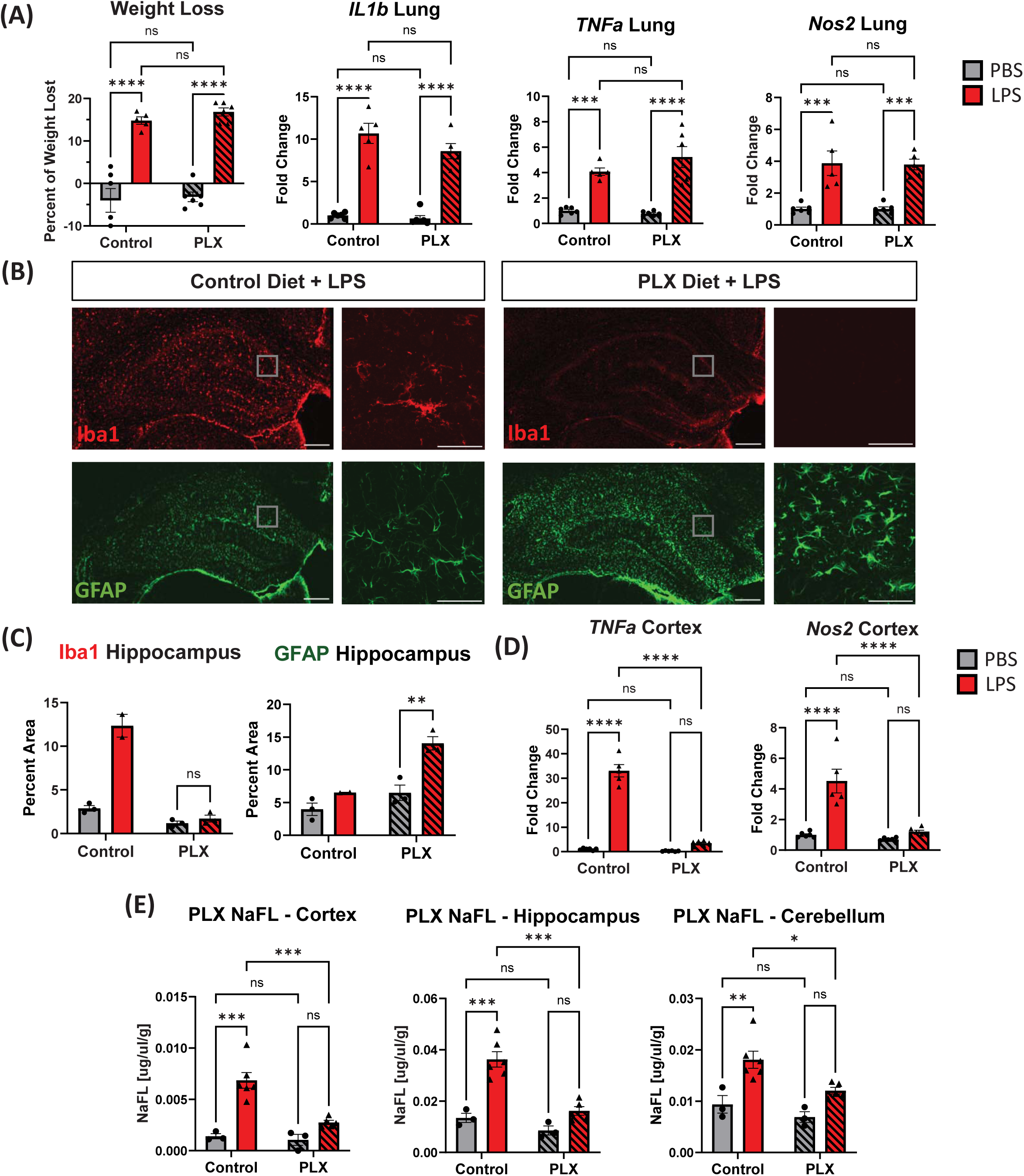
Microglia are required for LPS-induced inflammatory BBB breakdown at ZT13. (A) Peripheral cytokines and weight loss after 2 weeks of PLX3397 administration followed by 2d of PBS or LPS at ZT13. n = 5-6 mice per group. All graphs have significant main effect of LPS treatment only. (B) Representative images of hippocampal IBA1 and GFAP staining in control and PLX LPS-treated mice. Inset images are of the hippocampus at 40x magnification. (C) Quantification of IBA1 and GFAP hippocampal percent area. n = 2-3 mice per group. (D) LPS-induced cortical cytokine expression in PBS or LPS treated mice given control or PLX chow. n = 5-6 mice per group. (E) NaFL assay in PBS or LPS treated mice given either PLX or control chow. Scale bars: 200um in (B), instep is 50um.

We next sought to manipulate astrocyte activation to determine its role in neuroinflammation caused by evening LPS exposure. Since astrocytes are known to mitigate microglial reactivity after CNS-injury and are in close proximity to blood vessels, they are a prime candidate to serve as the first responders to LPS-induced peripheral cytokines (54). Activation of the JAK2-STAT3 pathway is a critical step in astrocyte activation in disease models (55). *In vitro*, LPS-stimulation induces an astrocytic STAT3-dependent release of TNF, leading to a loss of endothelial cell barrier properties (56). Thus, we chose to target the astrocyte STAT3 pathway in our model, utilizing a previously described AAV9 viral vector which expresses a constitutively active form of JAK2 (JAK2ca) behind a truncated GFAP promoter (57). Activated JAK2 phosphorylates STAT3, which then translocates to the nucleus and regulates the transcription of pro- and anti-inflammatory cytokines, including *Tnfa*, *Il1b*, and *Il6* (58). To ensure global viral spread, we performed intracerebroventricular injections of this viral vector, or an identical control vector expressing eGFP instead of JAK2ca, at postnatal day 0 and waited until the mice were 2-months-old before administering i.p. LPS for two days at ZT13. Mice treated with AAV-GFAP-JAK2ca showed increased hippocampal IBA1 and GFAP in response to LPS as compared to AAV-GFAP-eGFP treated animals, demonstrating that activating astrocyte JAK2-STAT3 prior to LPS exposure enhances both micro- and astrogliosis in the brain (Figure S3C). However, we observed no increase in BBB permeability at baseline or in response to LPS in the AAV-GFAP-JAK2ca animals. (Figure S3D). This finding suggests that while astrocytes respond to LPS independently of microglia, astrocyte activation alone is insufficient to induce BBB breakdown at ZT13 – at least through the JAK2/STAT3 pathway.

While microglia are not physically coupled to the NVU, they are still an intimate component of BBB function and increase perivascular localization during disease (Figure S3A) (16). Using the CSFR1 antagonist PLX3397, we found that microglia are required for evening LPS-induced BBB breakdown –despite persisting astrogliosis. While PLX3397 can exert off-target effects, including inhibition of Flt and C-kit, our observation that PLX3397 did not change LPS-induced inflammation in peripheral organs, but did impact the brain, strongly implicates microglia (59). It is still possible other CNS-resident CSFR1^+^ cells, such as perivascular macrophages (PVMs), play a role in LPS-induced inflammation. During many inflammatory diseases PVMs increase in number and vessel coverage; however, it is likely they are actually promoting vessel integrity, since their ablation increases leukocyte extravasation (60,61). Additionally, astrocyte activation in the absence of TLR4^+^ microglia indicates that astrocytes are responding to peripherally derived cytokines rather than parenchymal LPS – further supporting our peripheral cytokine data in Figure 1.

Our findings have several implications for human health. First, they might prompt investigation into diurnal variation in human BBB integrity in the setting of neuroinflammatory diseases. Second, the pathways that mediate this diurnal variation might be targeted therapeutically to prevent BBB breakdown in the setting of inflammatory diseases, particularly at times of peak vulnerability. Finally, diurnal susceptibility to inflammatory BBB permeability might be leveraged to allow better penetration of certain therapeutics, such as antibodies, into the brain. Chronotherapeutic regimens which incorporate information about diurnal BBB permeability dynamics may improve BBB penetration of anti-amyloid drugs, chemotherapies, and other agents. Our work, while a small first step, might open the door to such technologies.

## Methods

### SEX AS A BIOLOGICAL VARIABLE

This study examined LPS induced inflammation, for which there is a known sex-effect (62). However, we found no sex-effect on LPS-induced BBB breakdown in preliminary experiments, therefore we used exclusively male mice to reduce variability in the phenotype.

### BBB ASSAYS

100uL of 100mg/ml sodium fluorescein (Sigma-Aldrich: F6377) diluted in sterile PBS was injected subcutaneously, 45 minutes later mice were deeply anesthetized with i.p. pentobarbital (150mg/kg). 100uL of blood was collected by cardiac puncture before they were perfused with ice-cold PBS containing 3g/l heparin. Both hemispheres were dissected into regions and stored in pre-weighed tubes filled with 500uL PBS on ice. A small piece of peripheral and brain tissues were removed, flash frozen, and stored at -80°C for RNA analysis as described below. Tissue was weighed then homogenized for 3 minutes in a bullet blender and serum was added to a serum separator tube (BD Microtainer: 365967) before being spun down and diluted 1:200, tissue supernatant and serum were incubated 1:1 in 2% Trichloroacetic acid (Millipore Sigma: T9159) overnight (4°C). The samples were spun down again and diluted 1:1 in Borate Buffer (Honeywell: 33650) before being read on a plate reader along with a standard curve. Fluorescence per gram of CNS tissue was compared to fluorescence per uL of serum to determine the absolute amount of NaFL that crossed from the periphery to the brain for each mouse.

4uL/g body weight of 2% Evans Blue (Sigma-Aldrich, E2129) was injected retro-orbitally, 15 minutes later the mice were deeply anesthetized and perfused as described above. Whole brains were dissected out and imaged.

### CIRCADIAN RHYTHM DISRUPTION MODELS

For constant darkness experiments, mice were kept in standard 12:12 light:dark and after the lights turned off at 6pm they were kept off for the remainder of the experiment. For all experiments when mice were injected during the dark phase, red light was used and they were kept in darkness until fully anesthetized. Actigraphy was recorded using PIR infrared wireless sensors (Actimetrics) and circadian analysis was done using ClockLab Analysis software, version 6.1.02.

### DRUG ADMINISTRATION

LPS from *escherichia coli* O55:B5 (Sigma-Aldrich, L6529) was diluted to 2mg/ml in PBS and stored at -80°C. Immediately before each injection, LPS stock was thawed and diluted to 0.5mg/mL in PBS before being injected (i.p.) at a dose of 2mg/kg. Each LPS stock was not thawed more than twice before being discarded. To avoid batch effects, enough LPS was purchased at each time for an average of 3 experimental cohorts to be treated from the same stock. Mice were weighed each day prior to injection and the weight at the first day was used to determine dosing for the entire experiment.

### IMMUNOHISTOCHEMSITRY AND IMAGING

Immunohistochemical antibodies used in this study include: IBA1 (rabbit, Wako, 019-19741, 1:1000), GFAP conjugated to Alexafluor-647 (mouse, Cell Signaling Technologies, 3657S, 1:800), CD31 (rat, BD Pharmingen, 5500274, 1:250), and CLDN5 (mouse, Invitrogen, 35-2500, 1:100).

Mice were anesthetized and perfused as described above. For immunohistochemistry experiments, one hemisphere was dissected into regions, flash frozen, and stored at -80°C for RNA analysis as described below. The other hemisphere was post-fixed in 4% paraformaldehyde for 12 hours (4°C), then cryoprotected with 30% sucrose in PBS (4°C) for 24 hours. Brains were then sectioned on a freezing sliding microtome in 50-micron serial coronal sections and stored in cryoprotectant solution (30% ethylene glycol, 15% sucrose, 15% phosphate buffer in ddH20). Sections were washed in TBS x 3, blocked for 60 minutes in TBSX (TBS+ 0.4% Triton X-100) containing 3% donkey serum, and incubated overnight at 4°C in primary antibodies diluted in TBSX containing 1% donkey serum. Sections were then incubated for 1 hour at room temperature in TBSX with 1:1000 donkey fluorescent secondary antibody and mounted on slides using Fluoromount-G (Southern Biotech 0100-01) before coverslipping. For all analyses of tight junction and endothelial markers, mice were perfused with 4% PFA for 3 minutes, the whole brain was collected, and post-fixed in 4% paraformaldehyde for 4 hours (4°C) before moving to 30% sucrose for 24 hours (4°C). Brains were sectioned and stored as described above. Sections were blocked overnight in 1% Bovine Serum Albumin, 0.75% Triton x100, 5% donkey serum in PBS (4°C) before incubating in primary antibody diluted in PS/2 (0.5% BSA, 0.25% Triton x-100 in PBS) for 2 days (4°C). Sections were washed 3x in PS/2 then incubated for 2 hours at room temperature in PS/2 with 1:1000 donkey fluorescent secondary antibody, washed 2x in PS/2 and mounted as described above.

All fluorescent imaging was done on a Keyence BZ-X810 microscope. In general, laser intensity and exposure times were selected for each cohort of samples after a survey of the tissue, in order to select appropriate parameters that could then be held constant for all slides in that imaging session. These values varied by antibody, but all sections in a given cohort were imaged under identical conditions at the same magnification. For standard image analysis of epifluorescent images (such as determination of % area for antibodies such as anti-GFAP), TIFF image files were opened using ImageJ and converted to 8-bit greyscale files. Images with the dimmest and brightest intensity of staining, as well as some mid-range examples, were used to determine an appropriate threshold value that could optimally capture the intended staining across all conditions in that cohort, based on the judgment of the investigator. That threshold was then held constant across all images in the cohort, and black and white images of selected regions of interest were generated and quantified as % area stained using the Analyze Particles function. At least 2 adjacent sections per mouse per region were analyzed and averaged.

Perivascular gliosis analysis was done by taking 60x confocal images on a Zeiss LSM700 of blood vessels, the images were then exported and further analyzed using Imaris 10.0. In short, treatment groups were blinded to the investigator and 3D volumes for each channel were determined per image to account for LPS-effects on glial cell morphology, then an extended surface a distance of 8um away from blood vessels was applied. The volume of the channel colocalized to the extended surface compared to total channel volume as well as the volume of the 8um extended surface were measured. 40x images were taken with confocal resolution imaging and 2D maximum intensity projects were used for representative Images.

### PLEXIDARTINIB

Plexidartinib 3397 (PLX) was purchased from MedChemExpress (Hy-16749) and shipped to Research Diets, Inc (New Brunswick, NJ). PLX was formulated in AIN-76 diet at 400ppm and irradiated. PLX or control (AIN-76) chow were fed to mice ad libitum for 2 weeks prior to all PLX experiments.

### RNA QUANTIFICATION

Tissue was homogenized in 500ul of Trizol with beads added before running in a bullet blender for 3 minutes. TRIzol samples were then subjected to chloroform extraction (1:6 chloroform:TRIzol, followed by thorough mixing, and centrifugation at (12500 x g for 15 minutes). RNA was then extracted from the aqueous layer using the PureLink RNA Mini Kit according to manufacturer’s instructions. RNA concentration was measured on a Nanodrop spectrophotometer, then cDNA was made using a high-capacity RNA-cDNA reverse transcription kit (Applied Biosystems/Life Technologies) with 250ng-1 mg RNA per 20mL reaction. Real-time quantitative PCR was performed with ABI TaqMan primers and ABI PCR Master Mix buffer on ABI StepOnePLus or QuantStudio 12k thermocyclers. Taqman primers (Life Technologies) were used, and mRNA measurements were normalized to b-actin (Actb) mRNA for analysis. For larger experiments, microfluidic qPCR array measurements were performed by the Washington University Genome Technology Access Center using a Fluidigm Biomark HD system, again using Taqman primers. RNA sequencing was performed and values normalized as previously described (63). Volcano plots were made using Enhanced Volcano R package, heatmaps were made using the Pretty heatmap package, and X-Y plots were made using ggplot. Over-representation analysis dot plots of KEGG pathways were generated using the ClusterProfiler and enrichplot R packages.

### TRANSMISSION ELECTRON MICROSCOPY

Mice were anesthetized as described above and perfused with 10mL perfusion buffer (0.2mg/ml xylocaine and 20 units/ml heparin) then 20mL fixative solution (2.5% glutaraldehyde, 2% paraformaldehyde, and 0.15M cacodylate buffer with 2mM CaCl_2_), all kept at 37°C. Tissue was then post-fixed in ice-cold fixative solution overnight at 4°C. The brains were embedded in epoxy resin and the region of interest were cut into a 1mm x 1mm squares 70nm thick.

All imaging was done on a JEOL JEM-1400Plus Transmission Electron Microscope at 3-5,000x magnification. Capillaries were identified by morphology (<8um in diameter) with a circularity ratio <2. Treatment groups were blinded to the investigator and TIFF image files were opened using ImageJ and analyzed using the measure tool. The scale for each image was set depending on the magnification used. Diameter was determined as the distance across the vessel at the shortest point, luminal irregularity was the measured length of the plasma membrane divided by the estimated circumference using the elliptical tool, TJ length was the average of each measured TJ per vessel divided by the circumference, and vesicles were only counted if actively invaginating from the luminal side.

### VIRAL VECTORS

Jak2 constitutively active viral vector was made at the Hope Center Viral Vector core using Mammalian Gene Collection cDNA for mouse *Jak2* (Entrez ID 16452) generously gifted by Dr. Carole Escartin (CEA, Paris, France). Jak2ca cDNA was then packaged into an AAV9 envelope under expression of the truncated GFAabc1d promoter. A GFP tag was also included as a marker of Jak2ca expression. All viruses were administered via bilateral intracerebroventricular injection in newborn P0 pups as described previously (64). Two microliters of virus were injected at a concentration of 1.3x10^13^ gc/mL.

### STATISTICS

In all figures, graphs depict the mean + SEM, and N generally indicates the number of animals, unless otherwise noted in the figure legend. For confocal images, each datapoint depicts the mean of two sections with 6 technical replicates taken per section, each mouse is considered an N of 1. For TEM experiment, each datapoint depicts the mean for 28-32 technical replicates from one mouse, and each of these mean values is considered an N of 1. An F test was first performed for datasets with a single dependent variable and 2 groups, to determine if variances were significantly different. If not, 2-tailed unpaired T-test was performed. For datasets with 2 dependent variables, 2-way ANOVA was performed, and if main effect was significant, Tukey multiple comparisons test was then added for appropriate sets of variables. Outliers were identified using Grubbs test and were excluded. Statistical tests were performed with GraphPad Prism software, version 10.0.2. P values greater than 0.1 were noted as not significant (NS). P<0.05 was considered significant and was note with asterisks indicating the p-value: *P<0.05, **<0.01, ***<0.005, ****<0.00

### STUDY APPROVAL

All animal experiments were approved by the Washington University IACUC and were conducted in accordance with AALAC guidelines and under the supervision of the Washington University Department of Comparative Medicine. 7-week-old male C57/bl6J littermate mice were all obtained from Jackson Labs (Bar Harbor, ME) and allowed to acclimate for one week in our facilities before all experiments. For viral injection experiments embryonic day 18 timed-pregnant CD1 mice were ordered from Charles River (Wilmington, MA). Mice were housed in a 12-hour light/dark cycle (lights on at 6am and off at 6pm) and allowed ad libitum access to food and water unless otherwise stated during circadian rhythm disruption models.

## DATA AVAILABILITY

RNA-sequencing data submission to GEO is pending and accession number will be available upon publication. Values for all data points in graphs are reported in the Supporting Data Values file.

### Author Contributions

JHL: Designing research studies, conducting experiments, acquiring data, analyzing data, writing the manuscript

AP, MWK, CJN: conducting experiments

RD: Designing research studies

ESM: Designing research studies, overseeing research, providing reagents, writing the manuscript, providing funding.

## Acknowledgements

This work was funded by NIA grant R01AG054517 (ESM), National Science Foundation Grant DGE-2139839 and DGE-1745038 (JHL) and Washington University in St. Louis Brain Immunology and Glia (BiG) Center Interlab collaboration Grant (JHL). We thank Dr. Carole Escartin for the JAK2ca vector and Matthew Cain for advice on BBB assays. RNA sequencing was performed by the Genome Technology Access Center at the McDonnell Genome Institute at WashU. The Genome Technology Access Center (GTAC) is partially supported by NCI Cancer Center Support grant P30 CA91842 to Siteman Cancer Center, ICTS/CTSA grant# UL1TR002345 from the National Center for Research Resources (NCRR), a component of the National Institutes of Health (NIH), and NIH Roadmap for Medical Research. We thank the Washington University Center for Cellular Imaging (WUCCI) for assistance with confocal and electron microscopy. WUCCI is supported by Washington University School of Medicine, The Children’s Discovery Institute of Washington University and St. Louis Children’s Hospital (CDI-CORE-2015-505 and CDI-CORE-2019-813) and the Foundation for Barnes-Jewish Hospital (3770 and 4642).

