## Supplemental Data for "Microglia drive diurnal variation in susceptibility to inflammatory blood-brain barrier breakdown"

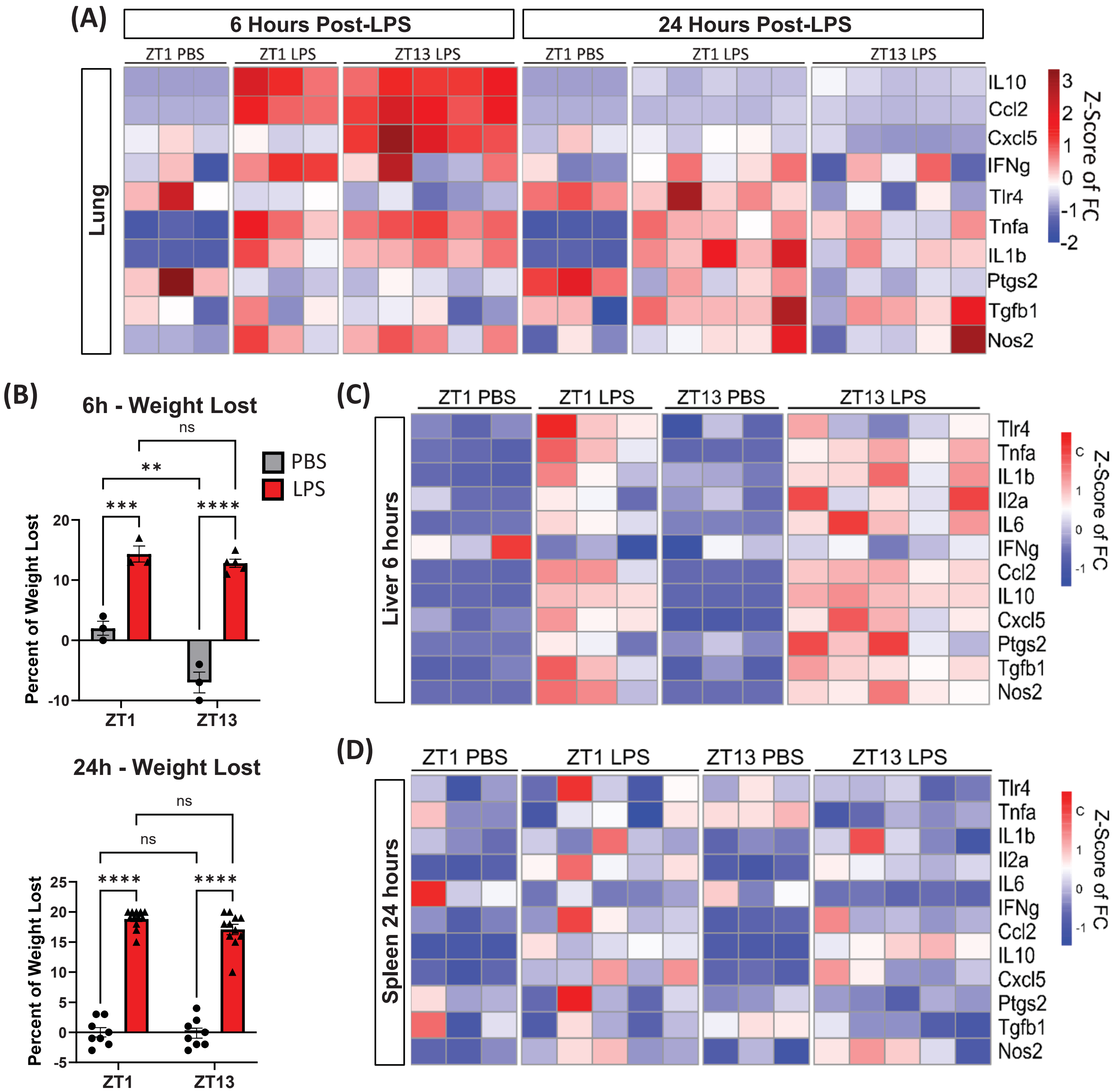


**Supplementary Figure 1: Absence of diurnal rhythm in peripheral immune activation at 6 or 24 hours post-LPS.** (A) Heatmap of selected gene expression in mouse lung tissue 6- vs 24- hours after final LPS injection at ZT1 or ZT12. Gene expression is normalized to ZT1 PBS 6 hpi group, Transcripts with significant TOD effect at 24 hpi are denoted with *. (B) Percent weight lost in PBS or LPS treated mice at ZT1 or ZT13 either 6 or 24 hours after LPS . (C) Heatmap of selected gene expression in mouse liver tissue 6 hours after final LPS injection at ZT1 or ZT12. Gene expression is normalized to ZT1 PBS group. (D) Heatmap of selected gene expression in mouse spleen tissue 24 hours after final LPS injection at ZT1 or ZT12. Gene expression is normalized to ZT1 PBS group.

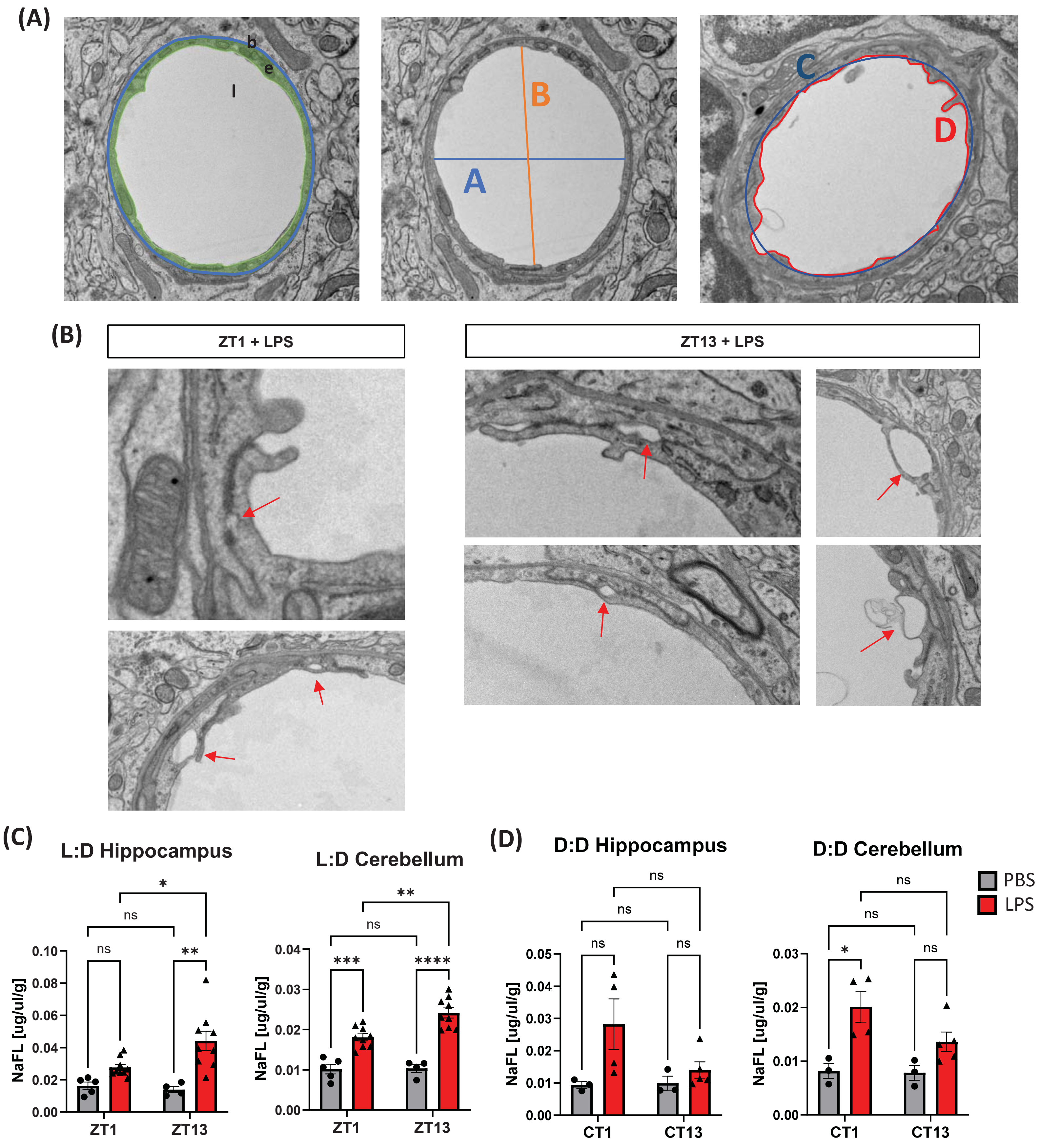


**Supplementary Figure 2: TJ disruption in select TEM mice and diurnal variation in BBB leak**

(A) Representative images of inclusion criteria for capillaries (b = basement membrane, e = endothelial cell, l = lumen); requirements for shortest diameter (A < 8 um) and vessel circularity (A/B < 2); luminal irregularity measurements (C = elliptical circumference, D = measured circumference). (B) Examples of TJ disruption and vacuole-like structures present in LPS treated mice at both ZT1 and ZT13 denoted by red arrow. (C) TOD differences in NaFL leak in the hippocampus and cortex of mice kept in 12h of light and dark. (D) TOD difference in NaFL leak in the hippocampus and cortex of mice kept in constant darkness (D:D) for 1d. n = 3-10 mice per group.

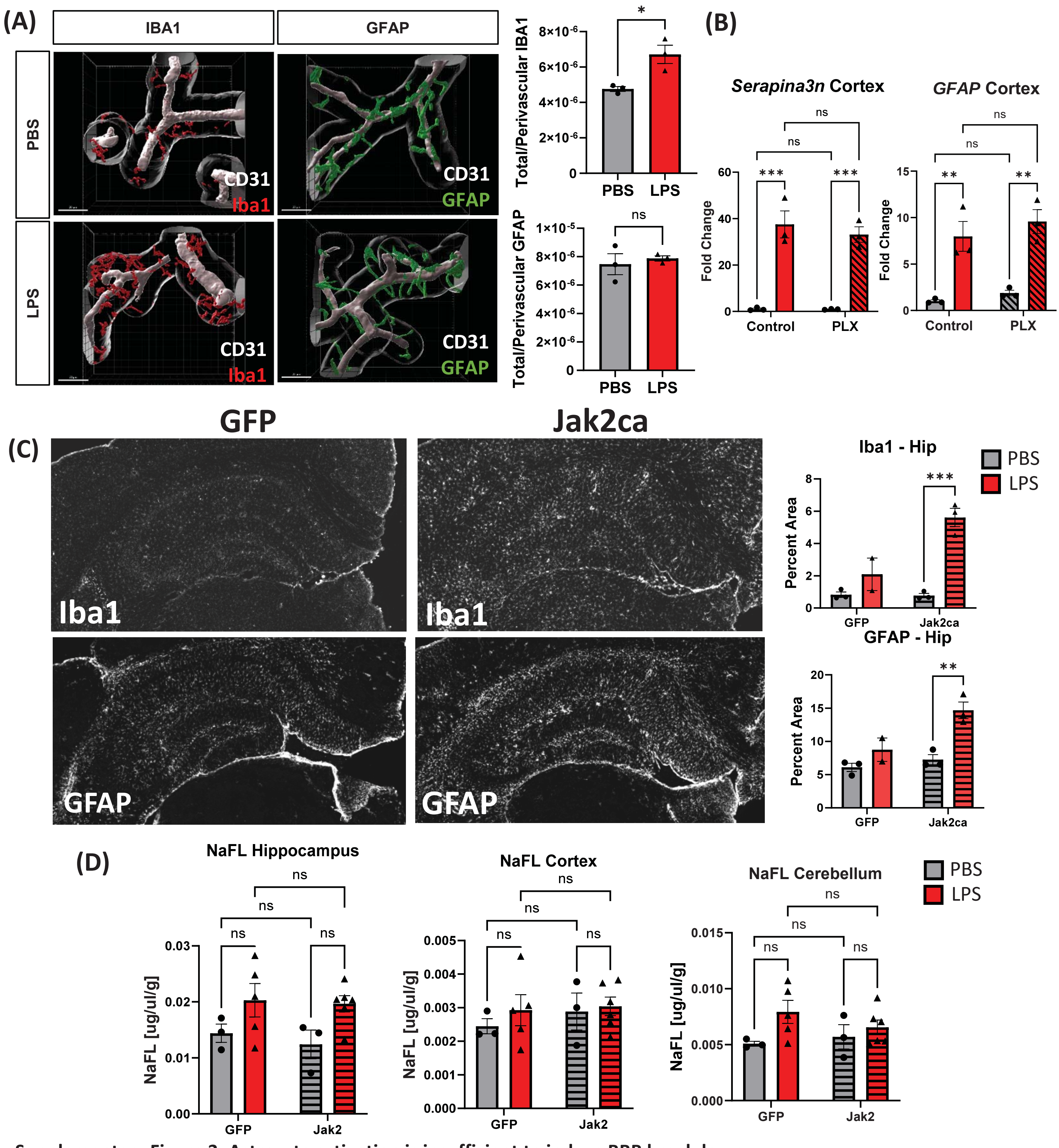


**Supplementary Figure 3: Astrocyte activation is insufficient to induce BBB breakdown**

(A) Volume of IBA1 and GFAP colocalized within 8 um of CD31, normalized to total volume, at ZT13 24 hpi. n = 3 mice per group; 3-6 vessels imaged per mouse. (A) *Serapina3n* and *Gfap* expression in PBS or LPS mice treated with 2 weeks of control or PLX chow. n = 3 mice per group. (B) NaFL assay in PBS or LPS treated mice given either PLX or control chow. n = 3-6 mice per group. (C) Representative images of IBA1 and GFAP in hippocampus of LPS treated GFP or Jak2ca injected mice, and quantification. n = 2-3 mice per group. (D) NaFL assay in PBS or LPS treated Jak2ca or GFP injected mice. n = 3-5 mice per group.
